# Proximal binding pocket Arg717 substitutions in *Escherichia coli* AcrB cause clinically relevant divergencies in resistance profiles

**DOI:** 10.1101/2021.12.16.473095

**Authors:** Martijn Zwama, Kunihiko Nishino

## Abstract

Multidrug resistance (MDR) in bacteria can be caused by the over-expression of multidrug efflux pumps belonging to the Resistance-Nodulation-Division (RND) superfamily of proteins. These intrinsic or acquired pumps can export a wide range of antibiotics. Recently, amino acid substitutions within these pumps have been observed in resistant clinical strains. Among others, two of these worrying gain-of-function mutations are R717L and R717Q in the proximal binding pocket of efflux pump AcrB (AcrB-Sa) found in azithromycin-resistant *Salmonella enterica* spp. We investigated the ramifications of these (and other) mutations in phylogenetically closely related AcrB from *Escherichia coli* (AcrB-Ec). We found that AcrB-Ec harboring Arg717 substitutions were significantly more effective in exporting all tested macrolides, with an up to 8-fold increase in the minimum inhibitory concentration (MIC) values (from 16 to 128 μg/mL for azithromycin). Interestingly, gain-of-function was also seen for fluoroquinolones (2-fold higher MICs), while there was a consistent loss-of-function for the export of novobiocin and β-lactam cloxacillin (2-fold lower MICs), pointing to a protein adaptation, which simultaneously partly compromised the efflux ability of other compounds with different molecular properties. Disk diffusion susceptibility testing corroborated the findings, as the R717Q and R717L mutant strains had significantly smaller inhibition zones for macrolides and fluoroquinolones and a larger inhibition zone for novobiocin, compared to the wild type. The spread and independent emergences of these potent efflux pump mutations highlight the necessity of control of, and adjustments to, treatments with antibiotics and the need for novel antibiotics and efflux pump inhibitors.

**Importance:** The over-expression of multidrug efflux pumps is a major cause of multidrug-resistant bacteria. The emergence of gain-of-function mutations in these pumps is additionally worrying, as these render last-resort antibiotic treatment options ineffective. The Arg717-mutations in related *Salmonella* Typhi and Paratyphi A AcrB caused azithromycin-resistant strains, with resistance levels past breakpoints. This is worrying, as azithromycin is a last-resort antibiotic after fluoroquinolones to treat typhoid and paratyphoid fever. Our findings that the same substitutions in closely related *Escherichia coli* AcrB cause an increase in MICs for both fluoroquinolones and macrolides is worrying as it shows the effect of one amino acid substitution on a multitude of treatment options. Additionally, the finding that the substitutions negatively impact the export of other drugs, such as β-lactam cloxacillin, suggests that treatment with multiple antibiotics may mitigate resistance and improve treatment. Our findings also imply the urge to develop novel antibiotics and inhibitors.

## Introduction

Antimicrobial Resistance (AMR) has become one of the major challenges in treating infectious diseases, as pathogenic microorganisms have become resistant to antibiotics (1, 2). Multidrug Resistance (MDR) is a worrying trend worldwide, as pathogens have been becoming non-susceptible to many or all available antibiotics (3, 4) and can be caused by reduced membrane permeability (5) or by over-expression of multidrug efflux pumps (5–7). Multidrug efflux pumps in clinical strains can be over-expressed from both intrinsic and acquired efflux pump genes (7–9), caused by mutations in their regulatory network (2). MDR in Gram-negative bacteria can be caused by the over-expression of efflux pump complexes belonging to the Resistance-Nodulation-Division (RND) superfamily (9, 10), which can expel a wide range of structurally unrelated antibiotics and toxic compounds from the cytoplasm and periplasm directly to the outside of the cells (11). Additionally, bacteria have further increased their resistance by adopting their efflux pumps by amino acid substitutions, limiting treatment options (12).

Among other gain-of-function mutations in RND-type efflux pumps from multiple pathogenic bacteria, specific amino acid substitutions in the AcrB efflux pump from *Salmonella enterica* Serovars Typhi and Paratyphi A have caused significant azithromycin-resistant phenotypes (12). These substitutions are present in the efflux pump’s proximal binding pocket (11–13) at the Channel 2 entrance (12–14). This is a concerning development, as in many countries, azithromycin is the last available and effective oral drug to treat typhoid (15, 16). In 2010, the first azithromycin-resistant *S.* Serovars Paratyphi A strains unresponsive to azithromycin treatment isolated from Pakistan were reported (17). *S.* Serovars Typhi and Paratyphi A strains isolated between 1995 and 2010 originating from Vietnam, Bangladesh, Cambodia, India, Laos, Nepal, and Thailand were analyzed in 2015 to determine azithromycin susceptibility breakpoints (>16 μg/ml for *S.* Typhi) (18). Hooda *et al*. (2019) first described the R717Q and R717L mutations present in *S.* Typhi and *S.* Paratyphi A clinical isolates, respectively, isolated between 2009 and 2016 from Bangladesh, with the first case dating at 2013. Specifically, they found 13 out of 1,082 strains with azithromycin resistance levels between 32–64 μg/ml, of which 12 harbored the R717Q/L mutation in the AcrB multidrug efflux pump (19). Similarly, in a recent study, *S.* Typhi isolates from Nepal from 2019 harboring the R717L mutation were non-susceptible to azithromycin (15). Genomic analysis of 133 *S.* Typhi strains collected in India between 1993 and 2016 showed that the two azithromycin non-susceptible strains contained the R717Q mutation in AcrB (20). Another recent study investigated an *S.* Typhi strain from Pakistan with an elevated azithromycin resistance level and also observed the R717Q mutation. Phylogenetic analysis showed the mutation likely spontaneously emerged, independently from the strains from Bangladesh (16). Furthermore, Sahib *et al.* (2021) investigated the emergence of azithromycin-resistant typhoidal *Salmonella* by analyzing 2,519 *S.* Typhi and Paratyphi A strains isolated in Bangladesh between 2016 and 2018. The 32 azithromycin-resistant *S.* Typhi strains all harbored the R717Q (29 strains) or R717L (3 strains) substitution. Five of the six other (slightly less) resistant *S.* Paratyphi A strains had the R717Q mutation. The first case of this substitution dated back to 2013, and they also predicted the Arg717-substitutions first emerged around 2010 and that a similar travel-related strain was isolated in the United Kingdom (21). Additionally, after cryo-EM and susceptibility analysis of a respective Arg714 substitution (R714G) in MtrD from *Neisseria gonorrhoeae* suggested a potential clinical relevance (22), three different substitutions (R714C/H/L) beside an overexpression-causing *mtrR* mutation were found in 12 out of 4,852 clinical isolates with elevated azithromycin resistance levels (23).

Here, we investigated the effect of the Arg717 substitutions R717R and R717L in *E. coli* AcrB (AcrB-Ec), which is phylogenetically closely related to *Salmonella* AcrB (AcrB-Sa). We investigated the effect of these mutations on the minimum inhibitory concentrations (MICs) of 20 different compounds, including planar aromatic cations (PACs), macrolides, β-lactams, quinolones, bile salts, and more. We also compared the effect of these two mutations to other interesting clinically relevant mutations, namely the proximal binding pocket mutations K823E/N (causing enhanced azithromycin resistance by MtrD) (22–25) and the distal binding pocket mutation G288D (causing fluoroquinolone resistance by AcrB-Sa in a *Salmonella* Typhimurium clinical strain) (26). Furthermore, we performed Kirby-Bauer disk diffusion susceptibility tests to determine the resistance significance for eight antimicrobials of interest (macrolides, fluoroquinolones, novobiocin, and minocycline).

## Results

To elucidate the impact of the Arg to Gln and Leu amino acid substitutions at position 717 (R717Q/L), we introduced the substitutions in *E. coli* AcrB (AcrB-Ec). The Arg717 amino acid is located at the Channel 2 (CH2) entrance of the proximal binding pocket (PBP) of the porter (drug efflux) domain of the RND efflux pump (Fig. 1a, red box). Arg717 is located close to a rifampicin molecule in the Access (or Loose (L)) monomer in one co-crystal structure (Fig. 1b,c) (13). This proximal pocket also holds a macrolide erythromycin molecule in another co-crystal structure, which entered the pocket through CH2 (Fig. 1b) (13, 14). *Salmonella enterica* (Typhi and Paratyphi A) AcrB and AcrB-Ec both comprise 1049 amino acids (Fig. S1) and share about 94% identity and 97% similarity (Fig. S2–4) (27). Both substitutions (R717Q and R717L) cause a decrease in side chain length, an increase in hydrophobicity, and a loss of a positive charge at the entrance of the PBP (Fig. 1c, S5), which we will discuss further in the Discussion section. Mutant and wild-type AcrB-Ec were expressed equally (Fig. S6).

**Figure 1.**
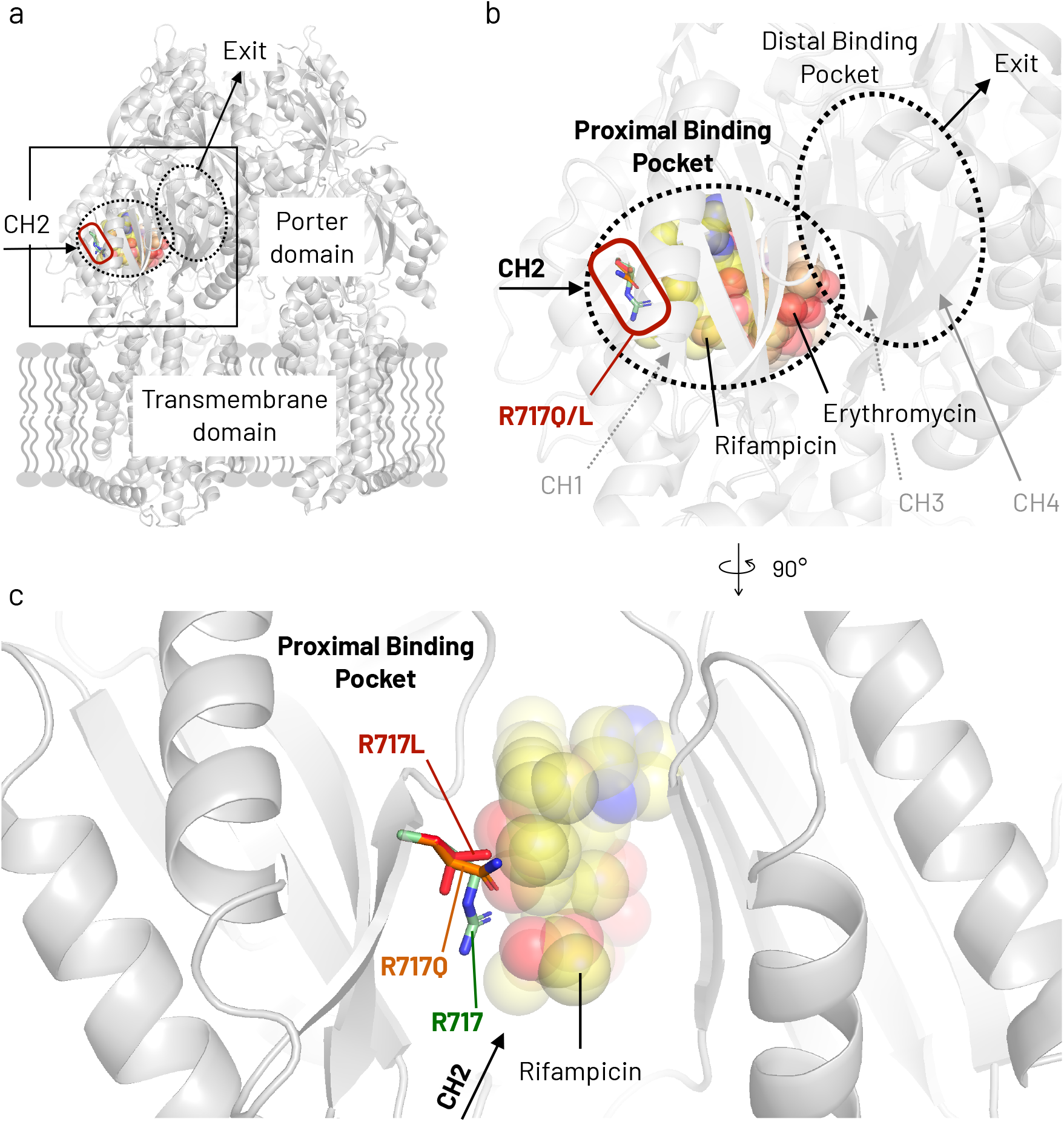
Location of Arg717 and the R717Q/L amino acid substitutions in AcrB-Ec. (A) Side-view of the entire AcrB-Ec trimer. (B) A close-up view of the rectangle area highlighted in panel A. Two drug molecules are present in the proximal binding pocket (rifampicin and erythromycin). The Arg717 residues and the R717Q/L substitutions are located at the entrance of the pocket. (C) Close-up inwards view of the proximal binding pocket and Arg717 area. Rifampicin can be seen behind the Arg717 residue and the R717Q/L substitutions. Abbreviations: CH1–4, Channel 1–4. Colors: red box, substitution area; yellow spheres, rifampicin; orange spheres, erythromycin; green sticks, Arg717; orange sticks, R717Q; red sticks, R717L. PDB accession codes: 4DX5 (for the main structure and the mutagenesis); 3AOC (for the erythromycin coordinates); 3AOD (for the rifampicin and the wild-type Arg717 side chain coordinates).

### R717Q and R717L substitutions cause divergent resistance spectra on solid agar plates

Table 1 shows the agar-plate (solid) minimum inhibition concentration (MIC) results of 20 different toxic compounds, including planar aromatic cations (PACs), macrolides, β-lactams, quinolones, bile salts, and other. Compared to wild-type AcrB-Ec, introducing the R717Q and R717L amino acid substitutions resulted in a divergent resistance spectrum. Both substitutions generally increased the MICs for macrolides and quinolones but decreased the MICs for novobiocin (NOV) and β-lactam cloxacillin (CLX). As for macrolides, the MICs increased most significantly for azithromycin (AZM) and clarithromycin (CLR). For CLR, wild-type AcrB-Ec expressing cells had an MIC of 128 μg/mL, while R717Q and R717L mutants had an MIC of > 256 μg/mL. As for AZM, the MIC for wild-type cells was 16 μg/mL, while for R717Q and R717L, the MICs were elevated to 64 and 128 μg/mL (a 4 and 8-fold resistance increase), respectively. MICs for erythromycin (ERY) were increased by 2-fold for both mutations compared to wild-type AcrB-Ec (from 128 to 256 μg/mL). Notably, for these viable R717Q/L cells, the growth seemed completely uninhibited, with healthy and thick colonies on the agar plates, even at their highest macrolide concentrations which showed growth (data not shown). Additionally, we observed a 2-fold increase in the MICs for certain fluoroquinolones, namely ofloxacin (OFX), levofloxacin (LVX), and moxifloxacin (MXF). Also, a significant 4-fold increase in MIC of monobactam aztreonam (ATM) was observed only for R717Q (0.125 μg/mL for mutant, and 0.03125 μg/mL for wild-type). Furthermore, only the R717L mutant increased the MIC for PAC ethidium bromide (EtBr) 2-fold. Interestingly, both R717Q and R717L mutants caused a 2-fold decrease in the MICs for NOV (128 μg/mL for the mutant, and 256 μg/mL for the wild type) and β-lactam CLX (also 128 μg/mL for the mutant, and 256 μg/mL for the wild type).

**Table 1.**
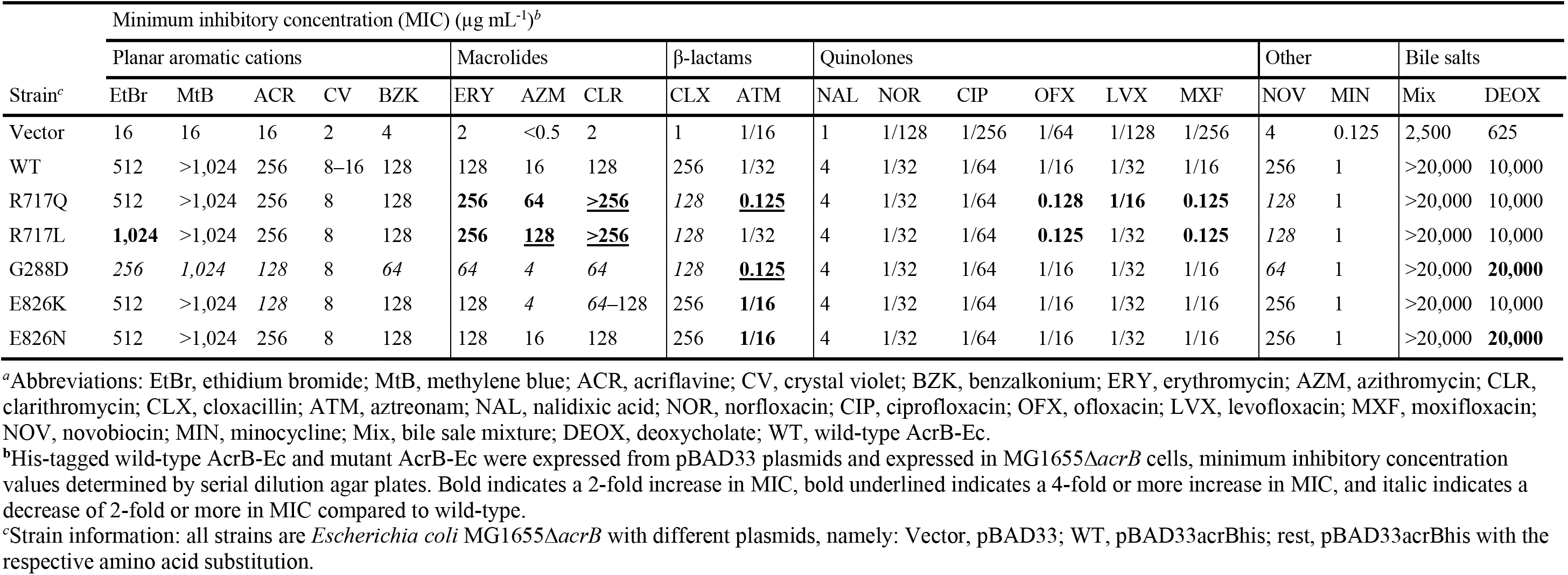
Antimicrobial susceptibility of AcrB-Ec expressing cells with several amino acid substitutions^*a*^

| Strain<sup>c</sup> | Minimum inhibitory concentration (MIC) (μg mL<sup>-1</sup>)<sup>b</sup> |  |  |  |  |  |  |  |  |  |  |  |  |  |  |  |  |  |  |  |
| --- | --- | --- | --- | --- | --- | --- | --- | --- | --- | --- | --- | --- | --- | --- | --- | --- | --- | --- | --- | --- |
|  | Planar aromatic cations |  |  |  |  | Macrolides |  |  | β-lactams |  | Quinolones |  |  |  |  |  | Other |  | Bile salts |  |
|  | EtBr | MtB | ACR | CV | BZK | ERY | AZM | CLR | CLX | ATM | NAL | NOR | CIP | OFX | LVX | MXF | NOV | MIN | Mix | DEOX |
| Vector | 16 | 16 | 16 | 2 | 4 | 2 | <0.5 | 2 | 1 | 1/16 | 1 | 1/128 | 1/256 | 1/64 | 1/128 | 1/256 | 4 | 0.125 | 2,500 | 625 |
| WT | 512 | >1,024 | 256 | 8–16 | 128 | 128 | 16 | 128 | 256 | 1/32 | 4 | 1/32 | 1/64 | 1/16 | 1/32 | 1/16 | 256 | 1 | >20,000 | 10,000 |
| R717Q | 512 | >1,024 | 256 | 8 | 128 | **256** | **64** | **>256** | *128* | **0.125** | 4 | 1/32 | 1/64 | **0.128** | **1/16** | **0.125** | *128* | 1 | >20,000 | 10,000 |
| R717L | **1,024** | >1,024 | 256 | 8 | 128 | **256** | **128** | **>256** | *128* | 1/32 | 4 | 1/32 | 1/64 | **0.125** | 1/32 | **0.125** | *128* | 1 | >20,000 | 10,000 |
| G288D | *256* | *1,024* | *128* | 8 | *64* | *64* | *4* | *64* | *128* | **0.125** | 4 | 1/32 | 1/64 | 1/16 | 1/32 | 1/16 | *64* | 1 | >20,000 | **20,000** |
| E826K | 512 | >1,024 | *128* | 8 | 128 | 128 | *4* | *64–128* | 256 | **1/16** | 4 | 1/32 | 1/64 | 1/16 | 1/32 | 1/16 | 256 | 1 | >20,000 | 10,000 |
| E826N | 512 | >1,024 | 256 | 8 | 128 | 128 | 16 | 128 | 256 | **1/16** | 4 | 1/32 | 1/64 | 1/16 | 1/32 | 1/16 | 256 | 1 | >20,000 | **20,000** |

<sup>a</sup>Abbreviations: EtBr, ethidium bromide; MtB, methylene blue; ACR, acriflavine; CV, crystal violet; BZK, benzalkonium; ERY, erythromycin; AZM, azithromycin; CLR, clarithromycin; CLX, cloxacillin; ATM, aztreonam; NAL, nalidixic acid; NOR, norfloxacin; CIP, ciprofloxacin; OFX, ofloxacin; LVX, levofloxacin; MXF, moxifloxacin; NOV, novobiocin; MIN, minocycline; Mix, bile salt mixture; DEOX, deoxycholate; WT, wild-type AcrB-Ec.
<sup>b</sup>His-tagged wild-type AcrB-Ec and mutant AcrB-Ec were expressed from pBAD33 plasmids and expressed in MG1655Δ*acrB* cells, minimum inhibitory concentration values determined by serial dilution agar plates. Bold indicates a 2-fold increase in MIC, bold underlined indicates a 4-fold or more increase in MIC, and italic indicates a decrease of 2-fold or more in MIC compared to wild-type.
<sup>c</sup>Strain information: all strains are *Escherichia coli* MG1655Δ*acrB* with different plasmids, namely: Vector, pBAD33; WT, pBAD33acrBhis; rest, pBAD33acrBhis with the respective amino acid substitution.

### Other clinical gain-of-function mutations generally cause increased susceptibility

To better understand the significance of the R717Q and R717L mutations at the channel 2 entrance of the PBP of AcrB-Ec and AcrB-Sa, we introduced other clinically relevant amino acid substitutions in AcrB-Ec, found in *Neisseria gonorrhoeae* MtrD and *Salmonella* Typhimurium AcrB-Sa. In MtrD, K823E and K823N were present in macrolide-resistant strains from Canada, the USA, and India (22–25). In AcrB-Sa, the G288D conferred increased resistance to fluoroquinolones in a clinical strain from the UK, causing a fatal infection (26). Therefore, we introduced the corresponding amino acid substitutions G288D, E826K, and E826N in AcrB-Ec. The location of Lys823 in MtrD corresponds to Glu826 in AcrB-Ec, and we were interested to see the influence of the MtrD substitution of Lys to Glu (K823E) as “reversed” substitution E826K in AcrB-Ec. Lys823 and Glu826 are located in the PBP of MtrD (22) and AcrB-Ec (13), respectively. These residues lie slightly deeper into the PBP, behind the Arg717 residue. On the other hand, Gly288 is located in the distal binding pocket (DBP) of AcrB-Ec (Fig. 1b) and AcrB-Sa, close to the hydrophobic trap (or inhibitor binding pit), where many substrates are found to be bound in the Binding (or Tight (T)) monomer in the crystal and cryo-EM structures of RND-type multidrug efflux pumps (13, 22, 28–33).

Table 1 shows the MIC results of G288D, E823K, and E823N mutant AcrB-Ec expressing cells. The substitution G288D in AcrB-Ec caused a decrease in the MICs for most PACs (EtBr, methylene blue (MtB), acriflavine (ACR), and benzalkonium (BZK)), all macrolides (ERY, AZM, and CLR), β-lactam CLX, and NOV. On the other hand, G288D in AcrB-Ec increased the MIC by 4-fold for monobactam aztreonam (ATM) and 2-fold for bile salt deoxycholate (DEOX). Furthermore, no increase in resistance was observed for fluoroquinolones as seen in the AcrB-Sa clinical strains, but also no decrease, as seen for PACs, macrolides, CLX, and NOV. E826K in AcrB-Ec caused an increased susceptibility for PAC ACR and macrolide AZM (and slightly for CLR); however, MICs were 2-fold higher for monobactam AZT. E826N caused only a 2-fold increase for ATM and DEOX. The increased susceptibility of AcrB-Ec(E826K) expressing cells to ACR, AZM, and CLR could correspond with the decreased susceptibility of macrolides AZM (23) and ERY (22, 24) for strains harboring the mutated MtrD(K823E).

### R717Q and R717L significantly increase growth ability in macrolide-supplemented liquid medium

The solid plate MIC results (Table 1) show an up to 8-fold increase in macrolide MICs for R717Q/L mutant AcrB-Ec expressing cells compared to wild-type cells. Furthermore, clinically relevant substitution G288D generally caused a decrease in the MICs by 2-fold. The reversed substitution E826K (corresponding to wild-type MtrD, compared the clinically observed K823E in MtrD) decreased the MICs for PACs, macrolides, CLX, and NOV by 2-fold. Hence, the 8-fold increase in MIC for AZM, the 2-fold for ERY, and the ≧ 4-fold for CLR seem to suggest a significant gain-of-function for AcrB-Ec regarding macrolide export ability. Therefore, we wanted to see the growth ability of AcrB-Ec(R717Q/L) expressing *E. coli* cells in macrolides, CLX and NOV supplemented liquid medium and compare it to wild-type *E. coli* MG1655 cells. Figure 2 shows the growth ability of wild-type, *acrB* knock-out, and mutant R717Q and R717L cells in macrolide supplemented broth. The top lane shows that wild-type cells have a significantly inhibited growth ability at 256 μg/mL ERY, and the growth is completely inhibited at 512 μg/mL. On the other hand, both R717Q and R717L have full growth ability at 256 μg/mL and still show significant growth ability under 512 μg/mL. Similarly, for CLR (middle lane), wild-type cells are basically completely inhibited at 256 μg/mL, while mutant cells can grow fully under the same concentration. At 512 μg/mL, mutant cells still have a slight growth ability. Furthermore, for AZM (bottom lane), wild-type cells already show a significant growth ability starting at 16 μg/mL and are completely inhibited at 32 μg/mL. On the other hand, mutant cells could fully grow even in 64 μg/mL AZM supplemented broth. Even in 128 μg/mL, the mutant strains seem to be ever so slightly viable (compared to the highest concentrations of ERY and CLR, Fig 2). These results corroborate the significant resistance increase caused by the R717Q/L substitutions for macrolides, especially relatively for AZM. Under all growth conditions (Fig. 2), the R717L AcrB-Ec expressing cells seemed to be somewhat more viable than R717Q AcrB-Ec expressing cells. Figure 3 shows the liquid growth of the same cells under CLX and NOV conditions. Plate MIC results (Table 1) indicated a 2-fold decrease in MIC for these compounds, and these results are reflected in Fig. 3, where R717Q/L AcrB-Ec expressing cells show an inhibited growth ability under various concentrations compared to wild-type AcrB-Ec expressing cells.

**Figure 2.**
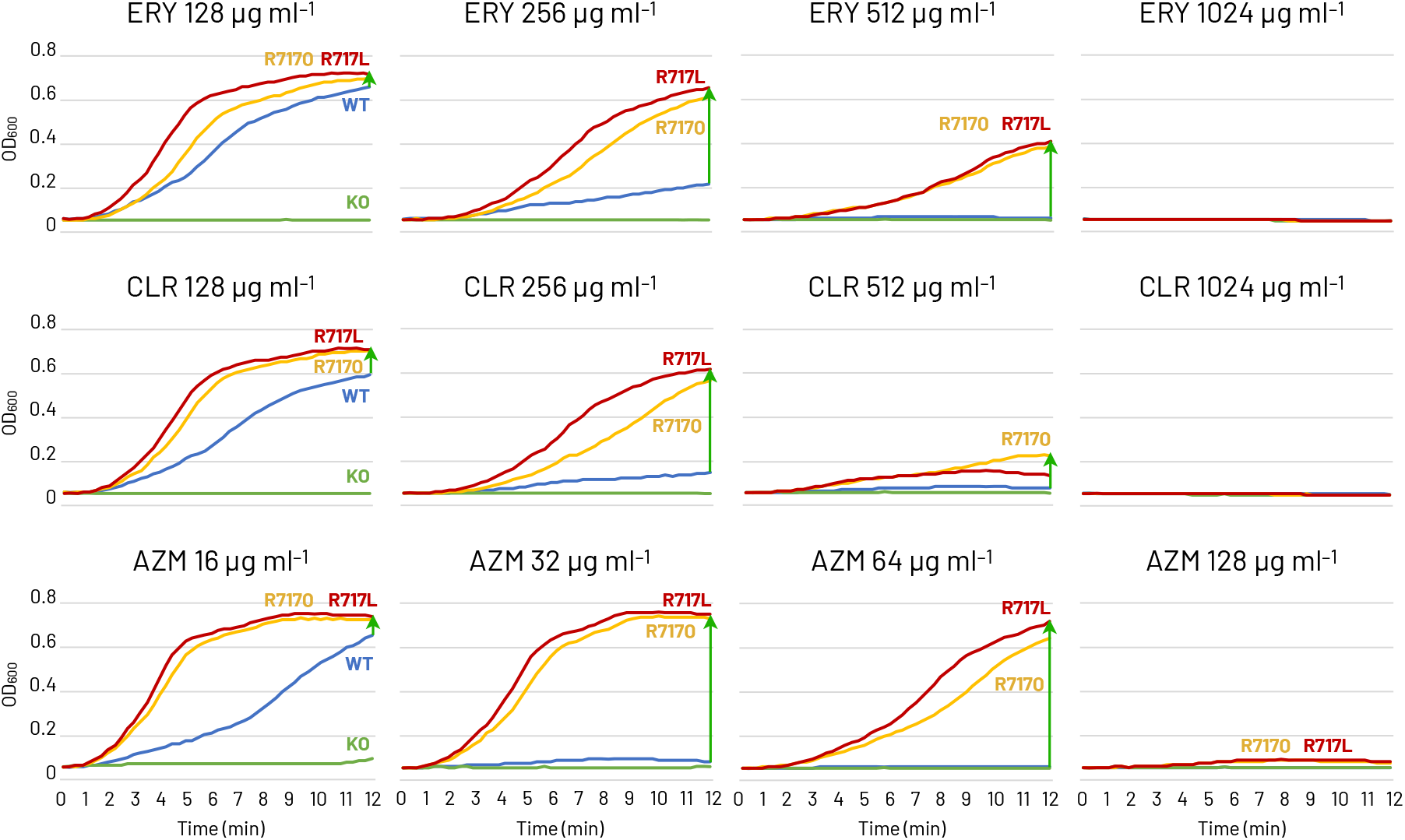
Growth ability of *E. coli* MG1655∆*acrB* expressing wild-type, R717Q, and R717L mutant AcrB under several macrolide concentrations. Top lane, growth ability under erythromycin; middle lane, under clarithromycin; bottom lane, under azithromycin. Colors blue, yellow, red, and green indicate wild-type AcrB, R717Q mutant, R717L mutant, and vector only, respectively. The green arrow mark indicates the increase in growth ability of the mutant strains compared to the wild-type strain. Abbreviations: ERY, erythromycin; CLR, clarithromycin; AZM, azithromycin. Experiments repeated twice providing similar results, shown is one of the results.

**Figure 3.**
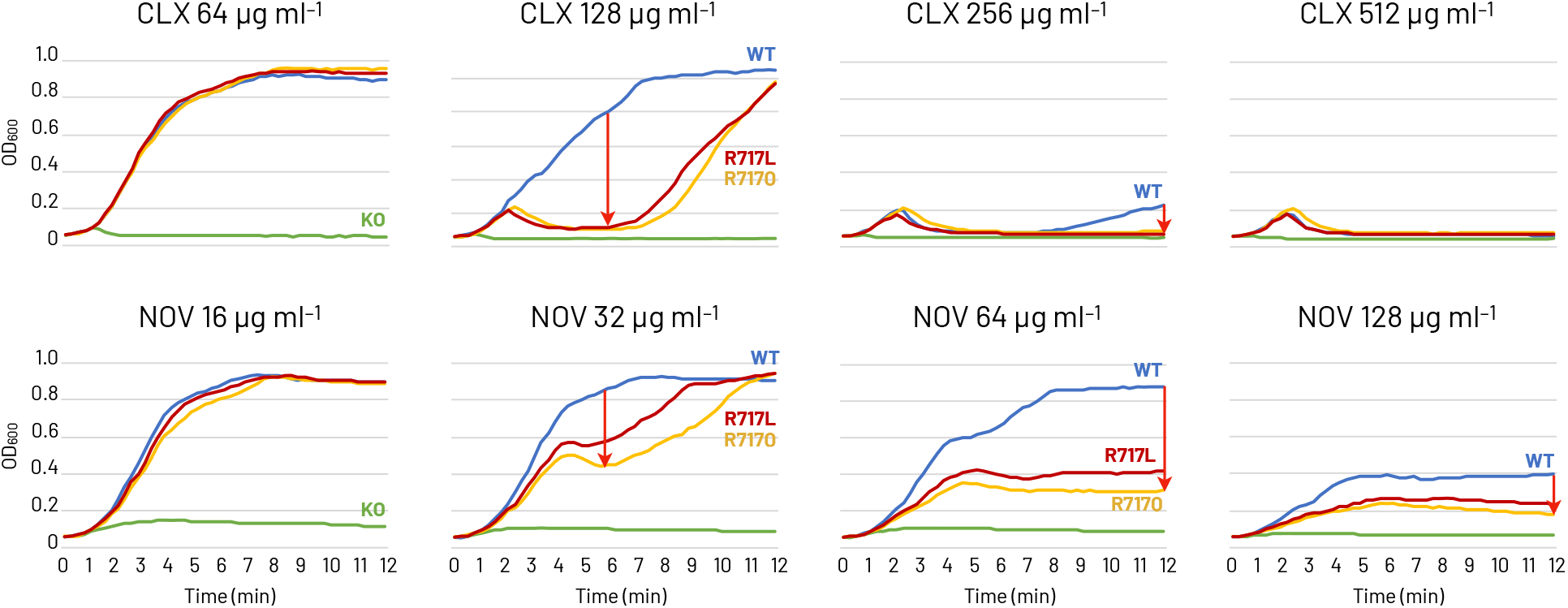
Growth ability of *E. coli* MG1655∆*acrB* expressing wild-type, R717Q, and R717L mutant AcrB under cloxacillin and novobiocin. Top lane, growth ability under cloxacillin; bottom lane, under novobiocin. Colors blue, yellow, red, and green indicate wild-type AcrB, R717Q mutant, R717L mutant, and vector only, respectively. The ref arrow mark indicates the decrease in growth ability of the mutant strains compared to the wild-type strain. Abbreviations: CLX, cloxacillin; NOV, novobiocin. Experiments repeated twice providing similar results, shown is one of the results.

### Kirby-Bauer disk diffusion susceptibility shows increased fluoroquinolone and macrolide resistance

To further check the clinical significance of the R717Q/L mutations, we performed Kirby-Bauer disk diffusion susceptibility tests. Besides a significant increase in resistance for macrolides, Table 1 shows that R717Q and R717L cause a mild but clinically significant 2-fold increase in the MIC for some fluoroquinolones (OFX, LVX, and MXF). In addition, recent studies on resistant *Salmonella* Typhi strains from Pakistan and Nepal observe isolates harboring the R717Q and R717L substitution in AcrB-Sa with reduced fluoroquinolone susceptibility; however, they note a double mutation in *gyrA* to contribute to this phenotype (15, 16). Nonetheless, we wanted to investigate further the consequences of the R717Q/L mutants in AcrB-Ec on fluoroquinolone resistance, as well as on macrolide resistance. Figure 4 shows the Kirby-Bauer disk diffusion susceptibility test results for wild-type, *acrB* knock-out, and mutants R717Q and R717L *E. coli* cells under macrolide (ERY, CLR, AZM) fluoroquinolone (OFX, LVX, MXF), NOV, and minocycline (MIN) supplemented growth conditions. When AcrB-Ec is expressed, the inhibition zones for all compounds decrease (Fig. 4, KO and WT). When R717Q and R717L mutant AcrB-Ec is expressed, the inhibition zones significantly further decrease for all three macrolides and all three fluoroquinolones (which can be seen by the naked eye in Fig. 4a and is quantified in Fig. 4b). It seems that R717L has a slightly more significant impact on the decrease of the inhibition zones than R717Q (Fig. 4b). Interestingly, R717L specifically decreases the inhibition zone for MIN slightly too. Similar to Table 1 and Figure 3, resistance to NOV is decreased when the R717Q/L mutations are introduced, with a decreased inhibition zone (Fig. 4a,b).

**Figure 4.**
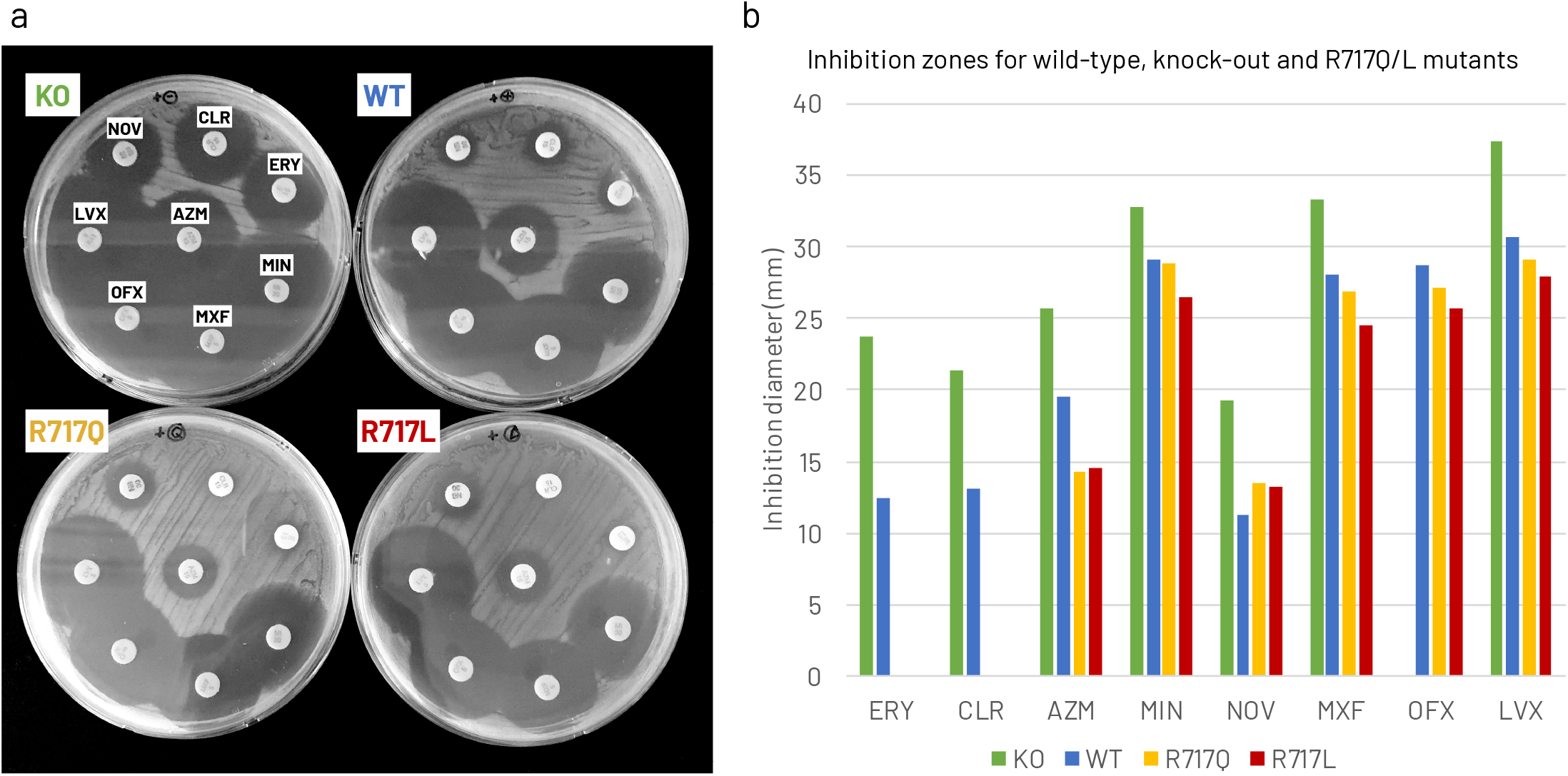
Kirby-Bauer disk diffusion susceptibility testing of *E. coli* MG1655∆*acrB* expressing wild-type, R717Q, and R717L mutant AcrB under various antibiotics. (A) Grayscale images of knock-out, wild-type, and mutant *E. coli* MG1655 grown on Mueller-Hilton agar plates, supplemented with resazurin and arabinose. Disks with specific antibiotics of interest can be seen on all four plates, along with the growth inhibition zones around the disks. (B) Quantification of the inhibition zones. The vertical axis shows the inhibition diameter in mm. Growth of the cells up to the antibiotic disk is denoted as 0 mm. (a–b) Colors blue, yellow, red, and green indicate wild-type AcrB, R717Q mutant, R717L mutant, and vector only, respectively. Abbreviations: KO, *acrB* knock-out (vector only); WT, wild-type, ERY, erythromycin; CLR, clarithromycin; AZM, azithromycin; NOV, novobiocin; LVX, levofloxacin; OFX, ofloxacin; MXF, moxifloxacin; MIN, minocycline.

**Figure 5.**
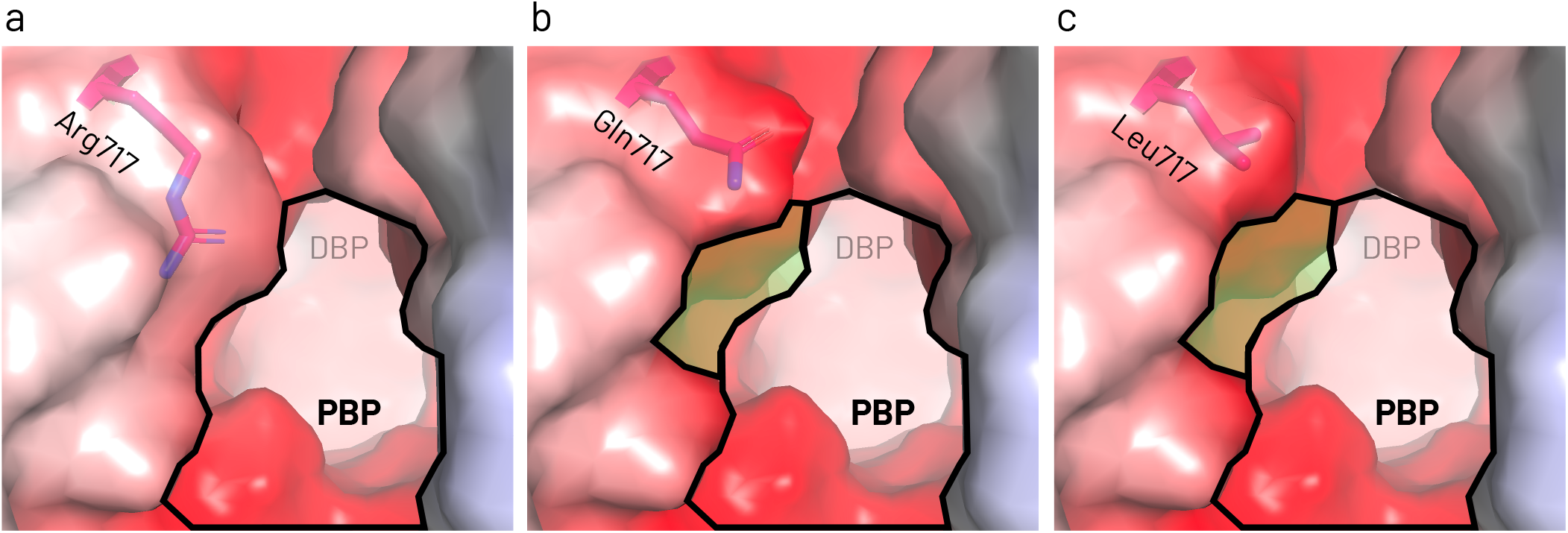
Comparison of the entrance area of the proximal binding pocket in the Access (or Loose (L)) monomer of AcrB-Ec for wild-type and the mutants. (A) Wild-type, (B) R717Q mutant, and (C) R717L mutant AcrB-Ec. (A–C) The residues of interest are depicted as pink sticks. The electrostatic surface is depicted from red to blue. The bold lines indicate the wild-type AcrB-Ec proximal binding pocket area at the Arg717 location. In (B) and (C), the green highlighted area depicts the increased area for R717Q (Gln717) and R717L (Leu717) mutant AcrB-Ec, respectively. These images show the entrance of the proximal binding pocket, leading to the distal binding pocket in the background. Abbreviations: PBP, proximal binding pocket; DBP, distal binding pocket. PDB accession code: 4DX5.

## Discussion

We showed that R717Q and R717L amino acid substitutions in AcrB-Ec from *E. coli* confers significantly increased resistance to macrolides, with an up to 8-fold increase in MIC. These findings corroborate the phenotypes of azithromycin-resistant *Salmonella* Typhi and Paratyphi A strains harboring the R717Q or R717L mutations in the AcrB-Sa efflux pump, found in multiple countries (including India, Nepal, Pakistan, and Bangladesh) (19–21). In addition, the same amino acid substitutions caused a 2-fold increase in the MICs of some fluoroquinolones, a phenotype also observed in *Salmonella* Typhi strains from Nepal (15) and Pakistan (16); however, in those studies, the phenotypes were attributed to mutations in *gyrA*. Interestingly, the R717Q and R717L substitutions caused a 2-fold reduction in the MICs for cloxacillin (a second-generation narrow-spectrum penicillin β-lactam) and novobiocin. This could (partly) explain the susceptibility of AcrB-Sa(R717Q)-harboring azithromycin-resistant *Salmonella* Typhi strains from Pakistan, which were still susceptible to third-generation cephalosporins (16). According to the EUCAST database, the theoretical azithromycin epidemiological cut-off MIC value for *E. coli* is 16 μg/mL, with only 28 strains found among 10,584 strains in literature with an azithromycin MIC of 32 μg/mL (besides 31 with 16 μg/mL, 907 with 8 μg/mL, and 9,618 with ≦ 4 μg/mL) (34). Our wild-type *E. coli* MG1655 strain has an azithromycin MIC of 16 μg/mL, and the R717Q and R717L mutants have an MIC of 64 and 128 μg/mL (Table 1), respectively, with fully healthy viable colonies growing up to these concentrations (data not shown). Liquid growth curves show the significant increase in viability of the mutant strains, with a “liquid MIC” of also 128 μg/mL for azithromycin, compared to 32 μg/mL for the wild-type strains (Fig. 2). Of the three tested macrolides (erythromycin, clarithromycin, and azithromycin), the mutants seemed to relatively increase the efflux ability of AcrB-Ec for azithromycin the most (4- and 8-fold increase in MICs for R717Q and R717L, respectively). In absolute terms, the MICs were increased most significantly for clarithromycin (> 256 μg/mL).

The amino acid substitutions R717Q and R717L (Fig. S5) cause a decrease in hydrophilicity by the removal of the positively charged residue. On the other hand, Gln is a polar residue, while Leu is a hydrophobic residue. Still, these two mutations both cause an increase in macrolide MIC. Macrolides are somewhat hydrophobic molecules, which also hardly dissolve in water, so the decrease in hydrophilicity may partly explain the increased drug efflux. Additionally, the length of the side chain is significantly reduced by both R717Q and R717L mutations. These substitutions both significantly enlarge the entrance of the proximal binding pocket (Fig. 1c, 5), which possibly also explains the enhanced efflux efficiency of macrolides by AcrB-Ec and AcrB-Sa, as macrolides are relatively large and high-molecular-mass-drugs (HMMDs) (13). Additionally, we previously showed that introducing bulky Trp residues at the lower entrance of the proximal binding pocket of AcrB-Ec (D566W and T678W) decreased the MICs significantly for erythromycin, in sharp contrast to ethidium bromide (14), corroborating the impact of space-limiting substitutions on the export of macrolides. The double effect of the increased space and the increased hydrophobicity could explain why the R717L mutant of AcrB-Ec seems slightly more active than R717Q (Table 1, Fig. 2). Furthermore, the inhibition zone of R717L AcrB-Ec was also significantly smaller than R717Q AcrB-Ec expressing cells for fluoroquinolones moxifloxacin, ofloxacin, and levofloxacin (Fig. 4). Interestingly, basically no difference was found between the two mutants on the growth under cloxacillin, while R717L seemed to have a higher growth ability than R717Q under novobiocin supplemented growth conditions (Fig. 3).

As explained before, besides a significant increase in macrolide resistance, and a relevant increase in fluoroquinolone resistance, we also observed a decrease in the MICs by 2-fold for cloxacillin and novobiocin. Therefore, it may be clinically interesting to combine multiple antibiotics to treat typhoid and paratyphoid fever to mitigate resistance and to enhance treatment. For example, a combination of β-lactams (including third-generation cephalosporins, as mentioned before) and azithromycin may hypothetically enhance the treatment of *Salmonella* infections. Additionally, we observed the significant gain-of-function in AcrB-Ec for the first time, showing that these mutations in other pathogenic bacteria may potentially have significantly worrying effects on clinical treatment options. These results further imply the importance of adjusted antibiotics treatments, as well as the need for novel antibiotics and efflux pump inhibitors.

## Materials and Methods

### Bacterial strains and growth conditions

The *E. coli* MG1655 strain (35) was used as wild-type strain, and ∆*acrB* (NKE96) (36) was derived from MG1655. Gene deletions were performed according to the method of Datsenko and Wanner, with recombination between short homologous DNA regions catalyzed by phage **λ**Red recombinase (37). The drug resistance markers were eliminated using plasmid pCP20 (37). The bacterial strains were grown at 37 °C in Luria-Bertani broth (38).

### Site-directed mutagenesis

A plasmid with the *acrB* gene (cloned from MG1655 *E. coli*, including a C-terminal hexahistidine-tag) pBAD33acrB was used for the site-directed mutagenesis. Point mutations were introduced by the PCR CloneEZ method (GenScript). Plasmids were transformed in *acrB-*deficient MG1655 *E. coli* cells.

### Drug susceptibility by MIC

Susceptibility testing in liquid and on solid media was determined by adding the toxic compounds by serial dilutions. Cell cultures of MG1655∆*acrB* harboring pBAD33, pBAD33acrBhis, or the mutated plasmids, were grown overnight and inoculated liquid cultures (supplemented with 10 mM arabinose) were incubated until OD_600 nm_ of around 0.6 and diluted to a final OD_600 nm_ of 0.05. For liquid growth curves, cell cultures were shaken at 37 °C, and OD_600 nm_ readings were performed by the Infinite M Nano (200 PRO series, Tecan). For LB agar MIC determination experiments, the diluted cells were stamped on LB agar plates (supplemented with 10 mM arabinose) and incubated at 37 °C overnight. MIC values were defined as the lowest drug concentrations at which the cells were no longer viable. AcrB expression levels (using 2 μg membrane fraction determined by a BCA assay) were analyzed by western blotting using an iBind western system (ThermoFisher), with an anti-his-tag mAb (MBL) as the primary and anti-mouse IgG HRP linked whole Ab from sheep (GE Healthcare) as the secondary antibody. AcrB-Ec and the R717Q, R717L, G288D, K823E, and K823N mutants were expressed in *E. coli* MG1655∆*acrB* cells in LB medium supplemented with 10 mM arabinose.

### Kirby-Bauer disk diffusion susceptibility test

Disk diffusion susceptibility was performed according to (39, 40), with slight modifications. In short: Mueller-Hinton agar plates were supplemented with resazurin to create colorimetric plates to make it easier to see the inhibition zones and cell viability by eye as resazurin reduction by viable cells change the color from blue to pink (41). *E. coli* MG1655∆*acrB* cells (harboring the previously mentioned four plasmids) were cultivated in LB broth overnight, and a dilution was grown until it reached an OD_600 nm_ of 0.5-0.6, and this was then diluted to OD_600 nm_ 0.1 with phosphate-buffered saline (PBS) buffer. Cells were streaked on pre-warmed plates with cotton swabs, and antibiotic disks (BD BBL Sensi-Disc from Becton, Dickinson and Company) were placed on each plate. Plates were then inverted and left overnight at 37 °C (42). The following day, inhibition zone diameters were measured with a digital ruler, and photos were taken.

## Data Availability

Data is available in the published article itself, and the supporting figures and tables are available as Supplementary Material. Other data that support the findings of this study are available from the corresponding authors upon request.

## Supplemental Material

Supplementary material is available at […].

FIG S1. PDF file, 89 kB

FIG S2. PDF file, 78 kB

FIG S3. PDF file, 82 kB

FIG S4. PDF file, 81 kB

FIG S5. PDF file, 58 kB

FIG S6. PDF file, 2.5 MB

## Acknowledgments

This research was supported by the Center of Innovation Program (COI) from the Japan Science and Technology Agency (JST), Grant-in-Aid for Scientific Research (Early-Career Scientists) (Kakenhi 20K16242), Grant-in-Aid for Scientific Research (Challenging Research (Exploratory)) (Kakenhi 18K19451) from Japan Society for the Promotion of Science (JSPS), CREST (JPMJCR20H9), the Takeda Science Foundation, the International Joint Research Promotion Program of Osaka University, and the Dynamic Alliance for Open Innovation Bridging Human, Environment and Materials from the Ministry of Education, Culture, Sports, Science and Technology-Japan (MEXT).

## References

1. World Health Organization. 2015. Global action plan on antimicrobial resistance. World Health Organization: Geneva, Switzerland. Available online: https://www.who.int/publications/i/item/9789241509763 (accessed on 01 December 2021).

2. Blair JMA, Webber MA, Baylay AJ, Ogbolu DO, Piddock LJV. 2015. Molecular mechanisms of antibiotic resistance. Nat Rev Microbiol 13:42–51.

3. World Health Organization. 2014. Antimicrobial resistance global report on surveillance: 2014 summary. World Health Organization: Geneva, Switzerland. Available online: https://apps.who.int/iris/handle/10665/112642 (accessed on 01 December 2021).

4. Walsh C. 2003. Where will new antibiotics come from? Nat Rev Microbiol 1:65–70.

5. Nikaido H. 2009. Multidrug Resistance in Bacteria. Annual Review of Biochemistry 78:119–146.

6. Li X-Z, Plésiat P, Nikaido H. 2015. The Challenge of Efflux-Mediated Antibiotic Resistance in Gram-Negative Bacteria. Clin Microbiol Rev 28:337–418.

7. Allen HK, Donato J, Wang HH, Cloud-Hansen KA, Davies J, Handelsman J. 2010. Call of the wild: antibiotic resistance genes in natural environments. Nat Rev Microbiol 8:251–259.

8. Levy SB, Marshall B. 2004. Antibacterial resistance worldwide: causes, challenges and responses. Nature medicine.

9. Blair JMA, Richmond GE, Piddock LJV. 2014. Multidrug efflux pumps in Gram-negative bacteria and their role in antibiotic resistance. Future microbiology 9:1165–1177.

10. Nikaido H. 1996. Multidrug efflux pumps of gram-negative bacteria. J Bacteriol 178:5853–5859.

11. Zwama M, Yamaguchi A. 2018. Molecular mechanisms of AcrB-mediated multidrug export. Res Microbiol 169:372–383.

12. Zwama M, Nishino K. 2021. Ever-Adapting RND Efflux Pumps in Gram-Negative Multidrug-Resistant Pathogens: A Race against Time. Antibiotics 10:774.

13. Nakashima R, Sakurai K, Yamasaki S, Nishino K, Yamaguchi A. 2011. Structures of the multidrug exporter AcrB reveal a proximal multisite drug-binding pocket. Nature 480:565–569.

14. Zwama M, Yamasaki S, Nakashima R, Sakurai K, Nishino K, Yamaguchi A. 2018. Multiple entry pathways within the efflux transporter AcrB contribute to multidrug recognition. Nat Commun 9:124.

15. Duy PT, Dongol S, Giri A, To NTN, Thanh HND, Quynh NPN, Trung PD, Thwaites GE, Basnyat B, Baker S, Rabaa MA, Karkey A. 2020. The emergence of azithromycin-resistant *Salmonella* Typhi in Nepal. JAC Antimicrob Resist 2:1–4.

16. Iqbal J, Dehraj IF, Carey ME, Dyson ZA, Garrett D, Seidman JC, Kabir F, Saha S, Baker S, Qamar FN. 2020. A Race against Time: Reduced Azithromycin Susceptibility in *Salmonella enterica* Serovar Typhi in Pakistan. mSphere 5:e00215–20.

17. Molloy A, Nair S, Cooke FJ, Wain J, Farrington M, Lehner PJ, Torok ME. 2010. First Report of *Salmonella enterica* Serotype Paratyphi A Azithromycin Resistance Leading to Treatment Failure. J Clin Microbiol 48:4655–4657.

18. Parry CM, Thieu NTV, Dolecek C, Karkey A, Gupta R, Turner P, Dance D, Maude RR, Ha V, Tran CN, Thi PL, Be BPV, Phi LTT, Ngoc RN, Ghose A, Dongol S, Campbell JI, Thanh DP, Thanh TH, Moore CE, Sona S, Gaind R, Deb M, Anh HV, Van SN, Tinh HT, Day NPJ, Dondorp A, Thwaites G, Faiz MA, Phetsouvanh R, Newton P, Basnyat B, Farrar JJ, Baker S. 2015. Clinically and Microbiologically Derived Azithromycin Susceptibility Breakpoints for *Salmonella enterica* Serovars Typhi and Paratyphi A. Antimicrob Agents Ch 59:2756–2764.

19. Hooda Y, Sajib MSI, Rahman H, Luby SP, Bondy-Denomy J, Santosham M, Andrews JR, Saha SK, Saha S. 2019. Molecular mechanism of azithromycin resistance among typhoidal *Salmonella* stains in Bangladesh identified through passive pediatric surveillance. Plos Neglect Trop D 13:e0007868.

20. Katiyar A, Sharma P, Dahiya S, Singh H, Kapil A, Kaur P. 2020. Genomic profiling of antimicrobial resistance genes in clinical isolates of *Salmonella* Typhi from patients infected with Typhoid fever in India. Sci Rep 10:8299.

21. Sajib MSI, Tanmoy AM, Hooda Y, Rahman H, Andrews JR, Garrett DO, Endtz HP, Saha SK, Saha S. 2021. Tracking the Emergence of Azithromycin Resistance in Multiple Genotypes of Typhoidal *Salmonella*. mBio 12:e03481–20.

22. Lyu M, Moseng MA, Reimche JL, Holley CL, Dhulipala V, Su C-C, Shafer WM, Yu EW. 2020. Cryo-EM Structures of a Gonococcal Multidrug Efflux Pump Illuminate a Mechanism of Drug Recognition and Resistance. mBio 11:e00996–20.

23. Ma KC, Mortimer TD, Grad YH. 2020. Efflux Pump Antibiotic Binding Site Mutations Are Associated with Azithromycin Nonsusceptibility in Clinical *Neisseria gonorrhoeae* Isolates. mBio 11:e01509–20.

24. Rouquette-Loughlin CE, Reimche JL, Balthazar JT, Dhulipala V, Gernert KM, Kersh EN, Pham CD, Pettus K, Abrams AJ, Trees DL, Cyr SS, Shafer WM. 2018. Mechanistic Basis for Decreased Antimicrobial Susceptibility in a Clinical Isolate of *Neisseria gonorrhoeae* Possessing a Mosaic-Like *mtr* Efflux Pump Locus. Mbio 9:e02281–18.

25. Wadsworth CB, Arnold BJ, Sater MRA, Grad YH. 2018. Azithromycin Resistance through Interspecific Acquisition of an Epistasis-Dependent Efflux Pump Component and Transcriptional Regulator in *Neisseria gonorrhoeae*. Mbio 9:e01419–18.

26. Blair JMA, Bavro VN, Ricci V, Modi N, Cacciotto P, Kleinekathӧfer U, Ruggerone P, Vargiu AV, Baylay AJ, Smith HE, Brandon Y, Galloway D, Piddock LJV. 2015. AcrB drug-binding pocket substitution confers clinically relevant resistance and altered substrate specificity. Proc National Acad Sci 112:3511–3516.

27. Zwama M, Yamaguchi A, Nishino K. 2019. Phylogenetic and functional characterisation of the *Haemophilus influenzae* multidrug efflux pump AcrB. Commun Biology 2:340.

28. Murakami S, Nakashima R, Yamashita E, Matsumoto T, Yamaguchi A. 2006. Crystal structures of a multidrug transporter reveal a functionally rotating mechanism. Nature 443:173–179.

29. Nakashima R, Sakurai K, Yamasaki S, Hayashi K, Nagata C, Hoshino K, Onodera Y, Nishino K, Yamaguchi A. 2013. Structural basis for the inhibition of bacterial multidrug exporters. Nature 500:102–106.

30. Morgan CE, Glaza P, Leus IV, Trinh A, Su C-C, Cui M, Zgurskaya HI, Yu EW. 2021. Cryoelectron Microscopy Structures of AdeB Illuminate Mechanisms of Simultaneous Binding and Exporting of Substrates. mBio 12.

31. Eicher T, Cha H, Seeger MA, Brandstätter L, El-Delik J, Bohnert JA, Kern WV, Verrey F, Grütter MG, Diederichs K, Pos KM. 2012. Transport of drugs by the multidrug transporter AcrB involves an access and a deep binding pocket that are separated by a switch-loop. Proc National Acad Sci 109:5687–5692.

32. Tam H-K, Foong WE, Oswald C, Herrmann A, Zeng H, Pos KM. 2021. Allosteric drug transport mechanism of multidrug transporter AcrB. Nat Commun 12:3889.

33. Ornik-Cha A, Wilhelm J, Kobylka J, Sjuts H, Vargiu AV, Malloci G, Reitz J, Seybert A, Frangakis AS, Pos KM. 2021. Structural and functional analysis of the promiscuous AcrB and AdeB efflux pumps suggests different drug binding mechanisms. Nat Commun 12:6919.

34. EUCAST. Antimicrobial wild type distributions of microorganisms. Available online: https://mic.eucast.org (accessed on 01 December 2021).

35. Blattner FR, Plunkett G, Bloch CA, Perna NT, Burland V, Riley M, Collado-Vides J, Glasner JD, Rode CK, Mayhew GF, Gregor J, Davis NW, Kirkpatrick HA, Goeden MA, Rose DJ, Mau B, Shao Y. 1997. The Complete Genome Sequence of *Escherichia coli* K-12. Science 277:1453–1462.

36. Senda Y, Yamaguchi A, Nishino K. 2008. The AraC-family regulator GadX enhances multidrug resistance in *Escherichia coli* by activating expression of *mdtEF* multidrug efflux genes. J Infect Chemother 14:23–29.

37. Datsenko KA, Wanner BL. 2000. One-step inactivation of chromosomal genes in Escherichia coli K-12 using PCR products. Proc National Acad Sci 97:6640–6645.

38. Sambrook J, Fritsch EF, Maniatis T. 1989. Molecular cloning: a laboratory manual (2nd ed.). Cold Spring Harbor, NY: Cold Spring Harbor Laboratory Press.

39. Bauer AW, Kirby WMM, Sherris JC, Turck M. 1966. Antibiotic Susceptibility Testing by a Standardized Single Disk Method. Am J Clin Pathol 45:493–496.

40. Kirby WM, Yoshihara GM, Sundsted KS, Warren JH. 1956. Clinical usefulness of a single disc method for antibiotic sensitivity testing. Antibiotics Annu 892–7.

41. Sener S, Acuner IC, Bek Y, Durupinar B. 2011. Colorimetric-Plate Method for Rapid Disk Diffusion Susceptibility Testing of *Escherichia coli*. J Clin Microbiol 49:1124–1127.

42. Hudzicki J. 2009. Kirby-Bauer Disk Diffusion Susceptibility Test Protocol. ASM 1–23.

